# Different issue types have different signal intensity on *b*=0 images and its implication on intravoxel incoherent motion (IVIM) analysis: examples of liver MRI

**DOI:** 10.1101/2021.03.11.431356

**Authors:** Ben-Heng Xiao, Yì Xiáng J. Wáng

## Abstract

Intravoxel incoherent motion (IVIM) theory in MRI was proposed to account for the effect of vessel/capillary perfusion on the aggregate diffusion weighted MR signal. The prevalent IVIM modeling is based on equation-1: SI_(b)_/SI_(0)_ = (1 -PF) × exp(-*b* × D_slow_) + PF × exp(-*b* × D_fast_) [1] where SI_(b)_ and SI_(0)_ denote the signal intensity of images acquired with the b-factor value of *b* and *b*=0 s/mm^2^, respectively. We recently reported that, for the liver and likely for other organs as well, IVIM modeling of the perfusion component is constrained by the diffusion component, with a reduced *D*_*slow*_ measure leading to artificially higher PF and *D*_*fast*_ measures. With higher b-value associated lower image signal of the targeted tissue, Euqation-1 is focused on describing the signal decay pattern along increasingly higher *b*-values by three IVIM parameters. Signal intensity at each *b*-value (i.e., SI_(b)_) is normalised by the signal intensity of *b*=0 image (i.e., SI_(0)_). We noted an apparent problem for Euqation-1. For example, if we want to compare the IVIM parameters of the normal liver parenchyma and a liver tumor, following Euqation-1 we will take the assumption that the SI_(0)_ of the normal parenchyma and the tumor are the same and considered equally as 1 (or 100) for the biexponential decay modelling. However, this assumption is invalid for many scenarios. From our liver IVIM database of 27 healthy female subjects, we chose six of the youngest subjects (20-27 yrs) and six of the oldest subjects (58-71 yrs) and measured the signals of the liver and left erector spinae muscle on *b*=0 and 2 s/mm^2^ images. The results show, while there was no apparent difference of left erector spinae muscle signal among the young and elderly groups, the elderly group’s liver SI_(0)_ is approximately 20 % lower than that of young group. This difference skewed the ratios of various SI_(b)_/SI_(0)_ and the followed IVIM parameter determination. The general trend is that lower liver SI_(0)_ is associated with lower *D*_*slow*_ and higher PF and *D*_*fast*_. If IVIM bi-exponential decay fitting starts from a very low non-zero b images (such as *b*=2 s/mm^2^ images), this problem persists. We performed an additional analysis of our IVIM database of five cirrhotic livers and the results show SI_(b=2)_ of cirrhotic right liver is positively associated *D*_*slow*_ (Pearson r=0.687), and negatively associated with PF (Pearson *r*=-0.733). Though the examples we used in this letter are on liver aging and liver fibrosis, the points discussed are expected to be generalisable to other pathologies and to other organs.

---

Intravoxel incoherent motion (IVIM) theory in MRI was proposed by Le Bihan et al to account for the effect of vessel/capillary perfusion on the aggregate diffusion weighted MR signal. The fast component of diffusion is related to micro-perfusion, whereas the slow component is linked to molecular diffusion. The prevalent IVIM modeling is based on Equation-1:

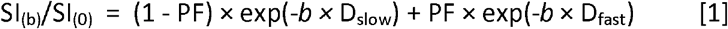

where SI_(b)_ and SI_(0)_ denote the signal intensity of images acquired with the *b*-factor value of *b* and *b*=0 s/mm^2^, respectively. Three parameters can be computed. D_slow_ (or *D*) is the diffusion coefficient representing the slow ‘pure’ molecular diffusion (unaffected by perfusion). The perfusion fraction (PF, or *f*) represents the fraction of the compartment related to (micro)circulation, which can be understood as the proportional ‘incoherently flowing fluid’ (i.e., blood) volume. D_fast_ (or *D*\*) is the perfusion-related diffusion coefficient representing the incoherent microcirculation within the voxel, which holds information for blood perfusion’s speed. We recently reported that, for the liver and likely for other organs as well, IVIM modeling of the perfusion component is constrained by the diffusion component, with a reduced *D*_*slow*_ measure leading to artificially higher PF and *D*_*fast*_ measures [1-3]. Thus, the perfusion component measure and the diffusion component measure cannot be separately determined.

The causes for this problem may be multiple. One of the causes could be that currently prevalent IVIM modeling (Euqation-1) does not fully consider the varied noise proportions of diffusion weighted images scanned under different acquisition conditions. During our analysis of liver IVIM data, we also noted another apparent issue for the Euqation-1. With higher *b*-value associated lower image signal of the targeted tissue, Euqation-1 is focused on describing the signal decay pattern along increasingly higher *b*-values by three IVIM parameters. Signal intensity at each *b*-value (i.e., SI_(b)_) is normalised by the signal intensity of *b*=0 image (i.e., SI_(0)_). Applying this analysis approach, for example if we want to compare the IVIM parameters of the normal liver parenchyma and a liver tumor, we will take the assumption that the SI_(0)_ of the normal parenchyma and the tumor are the same and considered equally as 1 (or 100) for the biexponential decay modelling. However, this assumption is invalid for many scenarios.

For our analysis of age and gender’s effect on liver IVIM measures, we noted that elderly female subjects had lower SI_(0)_ than that of younger females, which has also been described by Metens *et al* and may be related to elderly females’ liver have higher iron content [4]. As an example, if we use the women’s data from reference-1 and only choose those with clean and good quality, i.e., those had two good quality IVIM scans and we are able to use the mean values from these two scans, we will have 27 healthy women. From this pool of 27 subjects, we chose six of the youngest subjects (20-27 yrs, mean: 24.3 yrs) and six of the oldest subjects (58-71 yrs, mean: 63.3 yrs), and measured the signal of the right liver on *b*=0 and 2 s/mm^2^ images as described in reference-1, and also measured the signal of left erector spinae muscle on b=0 and 2 s/mm^2^ images. The results show, while there was no apparent difference of left erector spinae muscle signal among the young and elderly groups, the elderly group’ liver SI_(0)_ is approximately 20 % lower than that of young group (Fig-1). This difference skewed the ratios of various SI_(b)_/SI_(0)_ and the followed IVIM parameter determination. The general trend is that lower liver SI_(0)_ is associated with lower *D*_*slow*_ and higher PF and *D*_*fast*_ [1].

**Fig-1.**
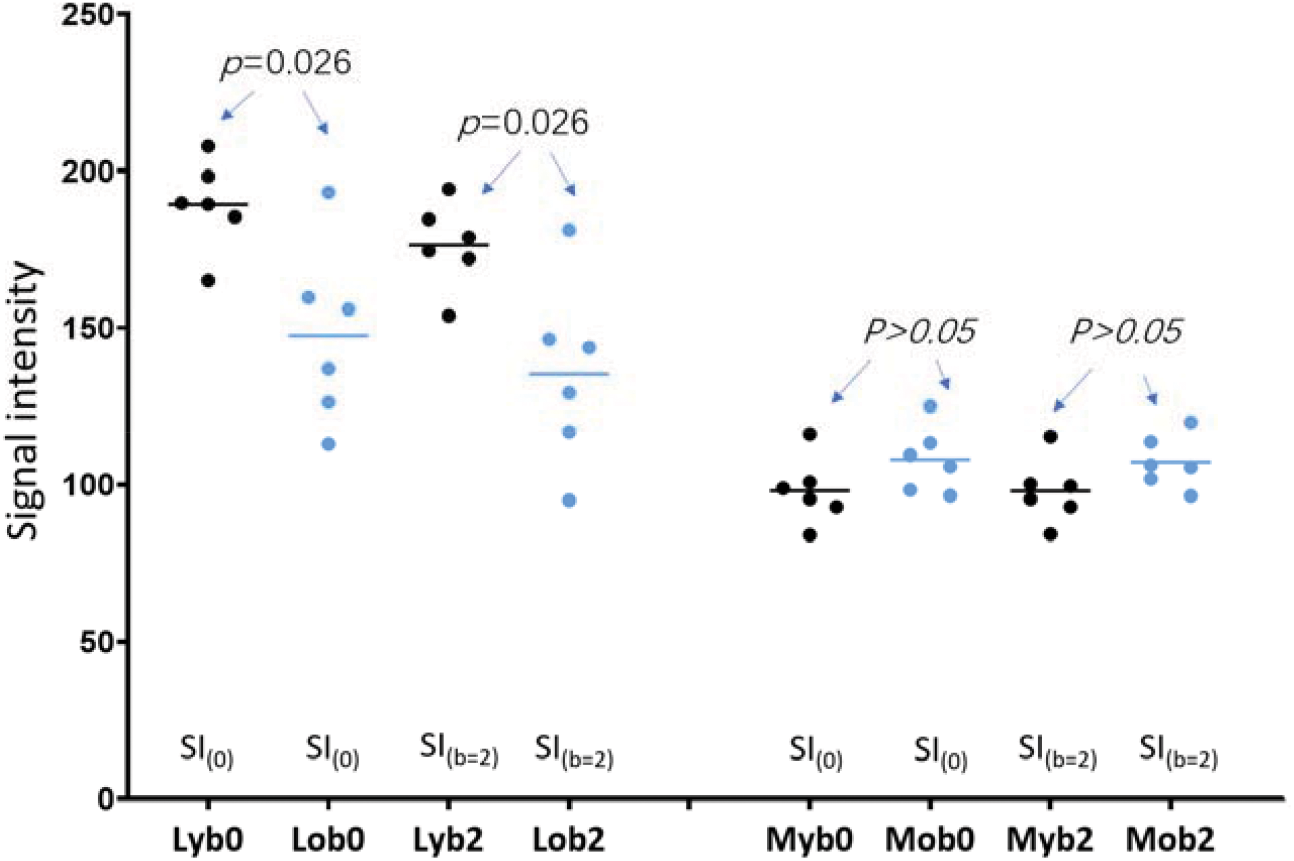
Absolute MR signal intensity (arbitrary unit/pixel) of right liver and left erector spinae muscle on *b*=0 (SI_(0)_) and 2 s/mm^2^ (SI_(b=2)_) images. The results of six young women and six elderly women are presented. Images were acquired on 1.5T. Lyb0: signal of young subjects’ liver on *b*=0 image. Lyb2: signal of young subjects’ liver on *b*=2 s/mm^2^ image. Lob0: signal of older subjects’ liver on *b*=0 image. Lob2: signal of older subjects’ liver on *b*=2 s/mm^2^ image. Mb0: signal of young subjects’ muscle on *b*=0 image. Myb2: signal of young subjects’ muscle on *b*=2 s/mm^2^ image. Mob0: signal of older subjects’ muscle on *b*=0 image. Mob2: signal of older subjects’ muscle on *b*=2 s/mm^2^ image.

There is currently a trend that *b*=0 images are not included for IVIM determination, instead IVIM bi-exponential decay fitting starts from a very low non-zero b images (such as *b*=2 s/mm^2^ images) [5-9]. This is to consider that, for perfusion rich tissues such as liver parenchyma, the IVIM signal decay from *b*=0 images actually does not follow a bi-exponential decay pattern (i.e., more likely follow a tri-exponential decay pattern) [10]. However, for IVIM analysis starting from a very low non-zero *b*-value images, the problem discussed above persists (Fig-1), as we have shown that lower liver SI_(b=2)_ is associated with lower *D*_*slow*_ and higher PF and *D*_*fast*_[1, 3]. We performed an analysis of five cirrhotic livers of the study of Li *et al* [5] and observed a very strong negative correlation between the measured *D*_*slow*_ and measured PF (Pearson *r*=-0.94) [3]. Our additional analysis (Fig-2) demonstrates SI_(b=2)_ of cirrhotic right liver is positively associated with *D*_*slow*_ (Pearson *r*=0.687), and negatively associated with PF (Pearson *r*=-0.733). This result is consistent with our observation of liver aging.

**Fig-2.**
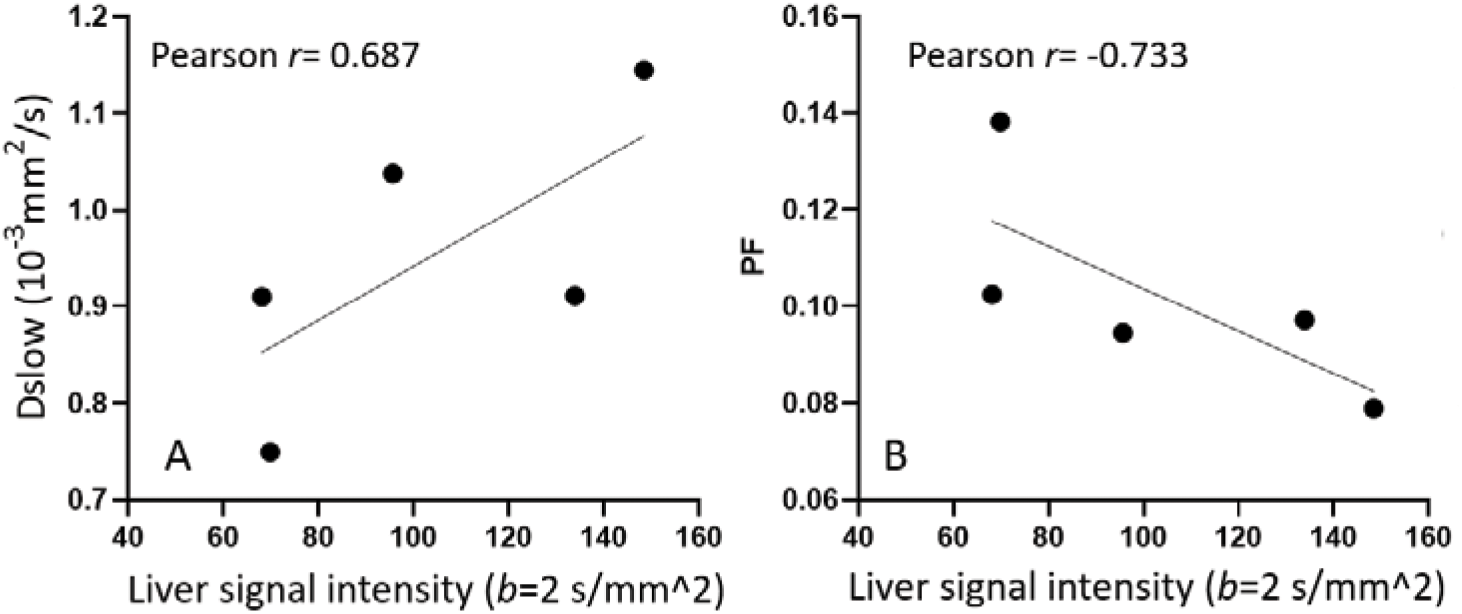
Absolute MR signal intensity (arbitrary unit/pixel) of liver on *b*=2 images and its correlation with *D*_*slow*_ and PF measure. Fiver cirrhotic livers’ images were acquired at 3T with 16 *b*-values of 0, 2, 5, 10, 15, 20, 25, 30, 40, 60, 80, 100, 150, 200, 400, and 600 s/mm^2^, analyzed by segmented fitting with threshold *b*-value of 60 s/mm^2^, with fitting starts from *b*=2 s/mm^2^ images (*b*=0 image excluded).

Though the examples we used in this letter are on liver aging and liver fibrosis, the points discussed are expected to be generalisable to other pathologies and to other organs.

